# Compulsion derived from incentive cocaine-seeking habits is associated with a downregulation of the dopamine transporter in striatal astrocytes

**DOI:** 10.1101/2024.11.19.624285

**Authors:** Maxime Fouyssac, Tristan Hynes, Aude Belin-Rauscent, Dhaval D. Joshi, David Belin

**Affiliations:** Department of Psychology, University of Cambridge, UK

**Author notes:** **Corresponding author:** Maxime Fouyssac or David Belin, Department of Psychology, University of Cambridge, Downing St., Cambridge CB2 3EB, UK, or.

**Keywords:** Astrocytes, Incentive habits, Cocaine, Dopamine Transporter, Compulsivity, Addiction

## Abstract

The development of compulsive cue-controlled -incentive- drug-seeking habits, a hallmark of substance use disorder, is predicated on an intrastriatal shift in the locus of control over behaviour from a nucleus accumbens (Nac) core - dorsomedial striatum network to a Nac core - anterior dorsolateral striatum (aDLS) network. Such shift parallels striatal adaptations to chronic drug, including cocaine self-administration, marked by dopamine transporter (DAT) alterations originating in the ventral striatum that spread eventually to encompass the aDLS. Having recently shown that heroin self-administration results in a pan-striatal reduction in astrocytic DAT that precedes the development of aDLS dopamine-dependent incentive heroin-seeking habits we tested the hypothesis that similar adaptations occurr following cocaine exposure. We compared DAT protein levels in whole tissue homogenates, and astrocytes cultured from ventral and dorsal striatal territories of drug naïve male Sprague Dawley rats to those of rats with a history of cocaine-taking or an aDLS dopamine-dependent incentive cocaine-seeking habit. Cocaine exposure resulted in a decrease in whole tissue and astrocytic DAT across all territories of the striatum. We further demonstrated that compulsive, i.e., punishment-resistant, incentive cocaine-seeking habits were associated with a reduction in DAT mRNA levels in the Nac shell, but not the Nac core-aDLS incentive habit system. Together with the recent evidence of heroin-induced downregulation of striatal astrocytic DAT, these findings suggest that alterations in astrocytic DAT may represent a common mechanism underlying the development of compulsive incentive drug-seeking habits across drug classes.

## Introduction

The relatively inflexible nature of drug-seeking that characterises cocaine use disorder (CUD) has been hypothesised to result from the development of compulsive incentive drug-seeking habits in vulnerable individuals (Belin *et al*., 2013). These incentive habits, which support drug foraging over long periods of time and promote compulsive relapse (Fouyssac *et al*., 2021a; Robbins *et al*., 2024), develop in rats over a long history of cue-controlled drug-seeking under second-order schedules of reinforcement (SOR) (Belin & Everitt, 2008; Murray *et al*., 2012; Murray *et al*., 2015). Under these conditions, which closely resemble drug foraging in humans (Koob, 2021), daily drug-seeking is invigorated and maintained over long periods of time by the response-contingent presentation of drug-paired cues acting as conditioned reinforcers (Everitt & Robbins, 2015). At the neural systems level, the development of incentive cocaine-seeking habits, which sets the stage for the transition to compulsive drug-seeking in vulnerable individuals (Jones *et al*., 2024), involves a progressive intrastriatal shift in the locus of control over behaviour. The acquisition of cue-controlled drug seeking depends on a basolateral amygdala - nucleus accumbens core (NacC) - posterior dorsomedial striatum (pDMS) network whereas well established, habitual cue-controlled drug seeking depends on a network involving the central amygdala, the NacC and anterior dorsolateral striatum (aDLS) dopamine-dependent mechanisms (Murray *et al*., 2012; Belin *et al*., 2013; Murray *et al*., 2015; Puaud *et al*., 2021). A similar progressive engagement of aDLS dopamine-dependent control over behaviour occurs following a protracted cue-controlled heroin seeking (Hodebourg *et al*., 2019), thereby suggesting that the development of incentive drug-seeking habits represents a core component of maladaptive drug-seeking behaviour across drugs.

Such recruitment of aDLS dopamine-dependent control over cue-controlled drug seeking likely depends on neurobiological adaptations within the striatum observed in response to exposure to drugs across species, such as alterations in glucose metabolism (Porrino *et al*., 2004) or alterations in the expression of the dopamine transporter (DAT) (Letchworth *et al*., 2001) and D2 Dopamine receptors (Moore *et al*., 1998), all originating in the ventral striatum and gradually spreading to the more lateral and dorsal territories of the striatum over the course of drug exposure. We recently demonstrated that striatal alterations in DAT protein levels over the course of heroin exposure occur within astrocytes (Hynes *et al*., 2024) and that a history of heroin self-administration results in a similar decrease in astrocytic DAT protein and mRNA levels throughout the striatum before the emergence of incentive heroin-seeking habits.

However, whether incentive cocaine-seeking habits and their compulsive manifestation are also associated with and preceded by a decrease in DAT expression in striatal astrocytes has not been elucidated. To this end, DAT protein levels were measured in total tissue μ-punches or cultured astrocytes harvested from ventral and dorsal striatal territories of male Sprague Dawley rats with a history of training under an incentive seeking habit-promoting SOR, a history of cocaine taking under continuous reinforcement, known to maintain pDMS-dependent goal-directed control over behaviour (Murray *et al*., 2012) or with no exposure to cocaine. In a second set of experiments, we examined whether the individual tendency to develop compulsive incentive cocaine-seeking habits was also associated with further alterations in DAT expression in striatal astrocytes. We previously demonstrated that heroin exposure resulted in similar downregulation of protein and mRNA levels in striatal astrocytes, but that only the latter remained reliably detected by qPCR (Hynes *et al*., 2024). Thus we assessed DAT mRNA levels from μ-punches of incentive habit- and compulsion-related striatal territories, namely the NacC, the aDLS (Belin *et al*., 2013; Giuliano *et al*., 2019; Jones *et al*., 2024) and the nucleus accumbens Shell (Chernoff *et al*., 2024), in rats that persevered in their incentive cocaine-seeking habits vs rats that readily relinquished their drug-seeking behaviour in the face of punishment (highly- (HC) vs non- (NC) compulsive rats, respectively) after a long history of training under a SOR (Jones *et al*., 2024).

## Methods and Materials

### Subjects

Our previous investigations on the role of astrocytic DAT in the development of incentive heroin seeking habits and all the findings related to the neurobehavioural basis of incentive cocaine seeking habits have been conducted exclusively on male rats. Therefore, in this study, we used 88 male Sprague Dawley rats (Charles River, UK), weighing 300-350g at the start of experiments and single-housed under a reversed 12h light/dark cycle (lights off at 7:00 AM). Rats were food restricted to reach 85% of their theoretical free-feeding body weight before behavioural training. Water was always available *ad libitum*. Experiments were conducted during the dark phase under the project license 70/8072 held by David Belin in accordance with the UK Animals (Scientific Procedures) Act 1986, amendment regulations 2012, following ethical review by the University of Cambridge Animal Welfare and Ethical Review Body (AWERB).

### Timeline

The experiments conducted in this study are schematically summarized in **Figure 1**.

**Figure 1:**
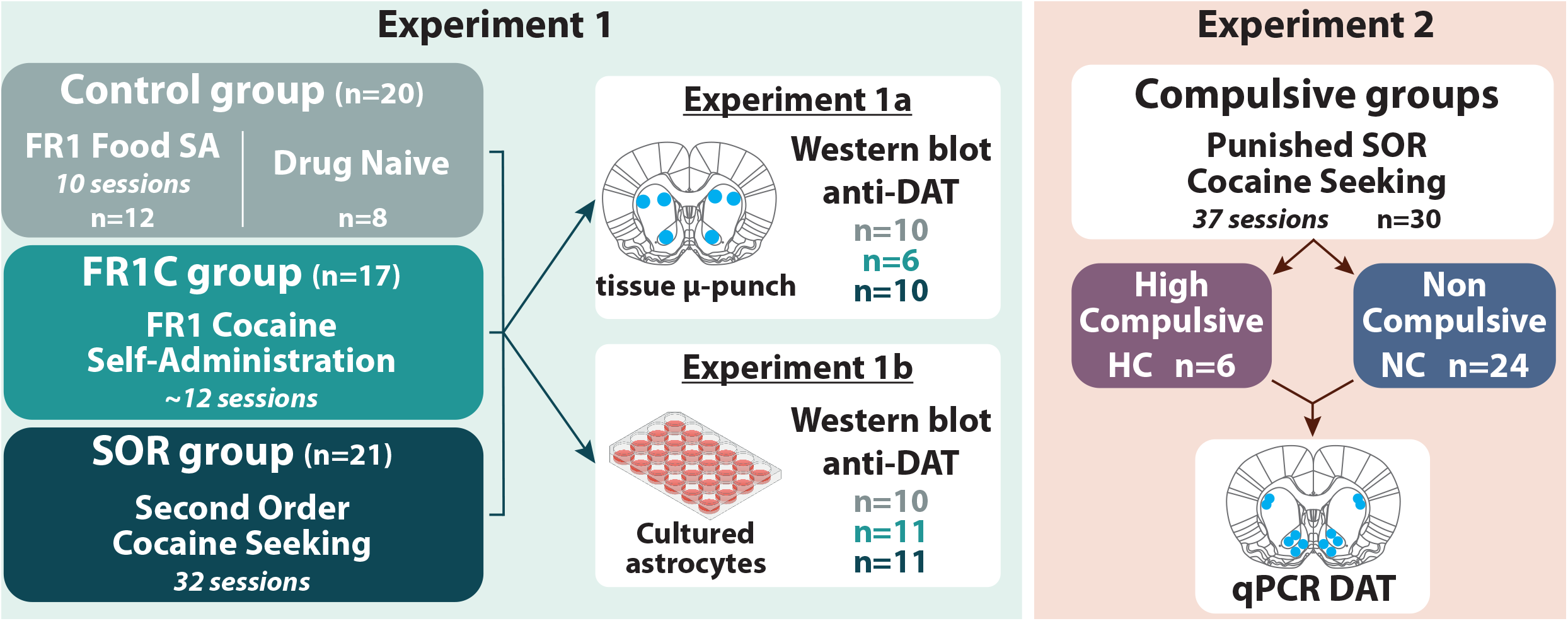
Experimental design. In **experiment 1**, rats (n=58) were trained instrumentally to respond under a fixed ratio 1 schedule of reinforcement either for food pellets (FR1 food group; n=12) or cocaine (FR1C group; n=17) or to seek cocaine under the control of the conditioned reinforcing properties of drug-paired CS, as operationalized under a FI15(FR10:S) second order schedule of reinforcement (SOR group; n=21). Other control animals were left undisturbed in their home cage (drug naïve group; n=8). In **experiment 1a**, we sought to characterise the impact of a history of drug taking or drug seeking on striatal DAT protein levels. The incentive drug seeking habit striatal system, namely NacC and aDLS, was investigated alongside the pDMS, pDLS and aDMS. Each structure was collected from the frozen brains of individuals of each of the four experimental groups, namely drug naïve untrained rats, FR1 food rats and cocaine exposed rats with (SOR rats) or without (FR1C rats) an incentive cocaine seeking habits. Total tissue DAT protein levels (including those expressed in striatal astrocyte and presynaptic dopaminergic neurons) were measured using western blot. To specifically investigate the influence of a history of cocaine exposure or well-established incentive cocaine seeking habits on the DAT protein levels of striatal astrocytes, we extracted primary astrocytes from freshly dissected striatal territories of individuals of each of these four groups in **experiment 1b**. DAT protein levels in these cultures were then assessed using western blot. Western blot data from the two control groups, FR1 food and drug naïve were pooled into a control group (CTL) since, as anticipated, they did not differ in their DAT protein levels. In **experiment 2**, rats (n=30) were trained to seek cocaine under a SOR until the compulsive nature of their incentive cocaine seeking habits was established over 3 sessions during which seeking responses were punished by mild electric foot shocks. Rats were characterised as high compulsive (HC; n=6) or non-compulsive (NC; n=24) by an unbiased k-means cluster analysis based on their resistance to punishment. Subsequently, astrocytic DAT mRNA levels were assessed by qPCR from micro-punches collected in the incentive habits striatal network, namely NacC and aDLS, as well as the NacS involved in a variety of compulsive behaviours.

### Drugs

Cocaine hydrochloride (Macfarlan-Smith, Edinburgh, UK) was dissolved in sterile physiological saline at a concentration of 2.5g/L.

### Surgery

All rats undergoing cocaine self-administration (n=68) were implanted with a home-made indwelling catheter into their right jugular vein under isoflurane anaesthesia as previously described (Jones *et al*., 2024). Perioperative analgesia was provided with Metacam (1 mg/kg, subcutaneously, Boehringer Ingelheim). Following surgery, rats received daily oral treatment with the analgesic for three days and an antibiotic (Baytril, 10 mg/kg, Bayer) for a week, which they first received on the day prior to surgery. Catheters were flushed with 0.1 ml of heparinized saline every other day after surgery and then before and after each daily self-administration session.

### Apparatus

All behavioural procedures were conducted in 24 operant chambers as previously described (Hynes *et al*., 2024). Scheduling and recording of experimental events were controlled by either MED-PC IV software (Med Associates, St. Albans, USA) or the Whisker software suite (Whisker, Cambridge, UK).

### Cocaine and Food Self-Administration

#### Fixed ratio 1 schedule of reinforcement for cocaine or food

Rats were trained instrumentally to respond for cocaine (0.25mg/infusion; 100μL/5 s) or food (one 45 mg food pellet) under a fixed ratio 1 (FR1) schedule of reinforcement as previously described (Hynes *et al*., 2024). Under this schedule, one active lever press resulted in the delivery of the outcome associated with the presentation of a conditioned stimulus (CS, cue light above the active lever). For cocaine self-administration, rats were trained for approximately 12 consecutive days, such that the number of cocaine infusions they received matched that of the individuals of the second order schedule of reinforcement group (SOR). For food reinforcement, rats were trained for ten daily sessions to control for instrumental and Pavlovian conditioning effects on DAT expression.

#### Cue-controlled cocaine seeking under a second-order schedule of reinforcement

Rats were trained to develop aDLS dopamine-dependent incentive cocaine seeking habits after extended exposure to a SOR as previously described (Fouyssac *et al*., 2021a). Chiefly, rats initially acquired cocaine self-administration under a FR1 schedule of reinforcement for three days. Rats were subsequently trained to seek cocaine under fixed interval schedules of reinforcement the duration of which increased across sessions from FI1 to FI2, FI4, FI8, FI10 and eventually FI15. After three days of training under a FI15 schedule of reinforcement, rats were further trained for twenty-one sessions under a FI15(FR10:S) second order schedule of reinforcement. Under these conditions, rats receive the drug upon responding after each 15min has elapsed. During these 15min seeking periods, rats receive a response-produced CS every 10^th^ lever press; thereby responding under the control of the reinforcing properties of the drug-paired cue.

#### Compulsive incentive cocaine seeking habits

Rats acquired cocaine self-administration under FR1 schedule of reinforcement for three days and were subsequently trained to seek cocaine under fixed interval schedules of reinforcement of increasing duration as described above. Following 17 daily sessions of SOR, rats were challenged with three daily sessions during which drug seeking was punished as previously described (Jones *et al*., 2024). During these three sessions, when rats were actively engaged in responding for the drug, namely during the last 7 minutes of each 15-minute interval, they received mild electric foot-shocks (1 sec duration, 0.45 mA) every 16^th^ lever press. This contingency enables the suppression of responding in resilient animals while minimising the probability of counterconditioning and avoiding extinction of drug seeking behaviour. Each shock was paired with a cue light located on the top middle region of the wall distinct from that paired with cocaine infusions.

Following the punished sessions, rats were re-exposed to three baseline SOR sessions. An unbiased K-means cluster analysis *(Giuliano et al., 2021; Marti-Prats et al., 2021; Jones et al., 2024)* was carried out on the number of shocks rats were willing to receive to maintain their incentive habits during the first, drug-free, interval of the last two punishment sessions, when a significant and stable suppression of seeking was observed in non-compulsive individuals, as shown in **Figure 4**.

To ensure that potential differences in resistance to punishment were not attributable to a differential sensitivity to nociceptive stimuli, rats were tested in a hot plate test before the punishment sessions. As previously described *(Fouyssac et al., 2021b)*, rats were placed on a hot plate calibrated to remain at a stable temperature of 52ºC. The time elapsed before the appearance of pain-associated behaviours, including paw-licking and jumping, was measured and considered a direct indicator of pain threshold *(Woolfe & Macdonald, 1944)*.

### Astrocytes Primary Cell Cultures

Astrocytes primary cultures from the Nucleus Accumbens Core (NacC), the anterior and posterior dorsomedial striatum (aDMS and pDMS, respectively) and the anterior and posterior dorsolateral striatum (aDLS and pDLS, respectively) were carried out under the exact same conditions as previously described (Hynes *et al*., 2024).

### Immunohistochemistry

Immunohistochemistry assays were carried out under the exact same conditions as previously described (Hynes *et al*., 2024). Cells were incubated overnight with a rabbit anti-GFAP primary (1:500; Merck Millipore, #04-1062), a mouse anti-Iba-1 primary (1:500; Merk Millipore, #MABN92), a chicken anti-NeuN primary (1:1000; Merc Millipore #ABN91), or a mouse anti-GFAP primary (1:500; Merck Millipore #MAB360). Secondary antibodies were incubated for 90 minutes at room temperature: anti-rabbit Alexa Fluor® 647 (1:800; Merck Millipore, #AP187SA6), anti-mouse Alexa Fluor® 488 (1:800; Merck Millipore, #AP124JA4), anti-mouse Alexa Fluor® 568 (1:1000; Invitrogen #A-11004), or anti-chicken Alexa Fluor® 647 (1:1000; Invitrogen #A-21449). Cells were then treated with DAPI (Life Technology, #D1306) and mounted on slides with DPX mounting medium (Fischer, #D/5319/05).

### Western blot

Protein extraction, electrophoresis, antibody incubation, visualisation and quantification were carried out in the exact same conditions as those used for experiments 1 and 2 of our recently published study (Hynes *et al*., 2024). Raw and uncropped western images used in **Figures 2 and 3** are presented in the supplement **Figure S1**.

**Figure 2:**
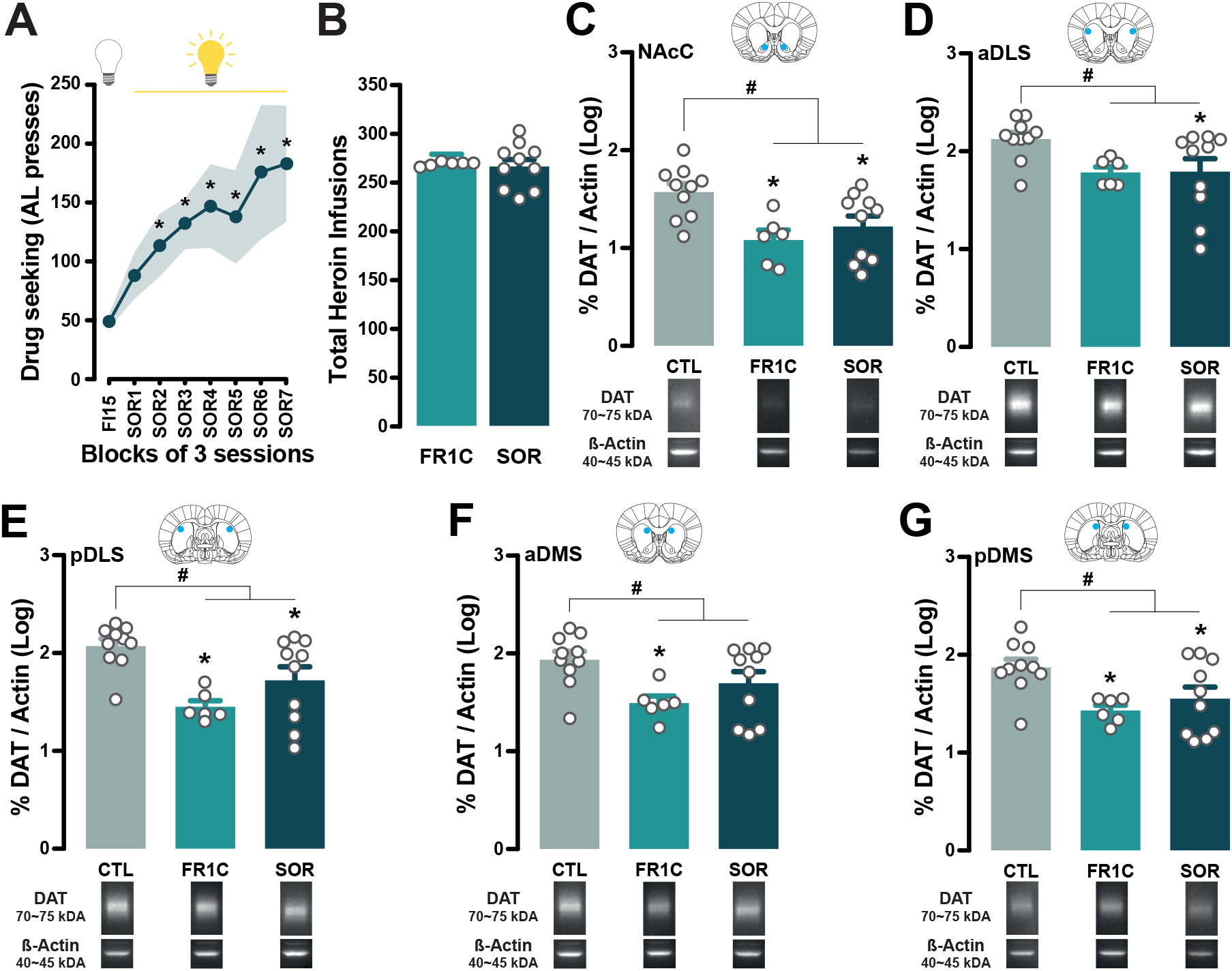
Cocaine exposure triggers a downregulation of DAT protein levels across the striatum. **(A)** rats trained to seek cocaine under a second-order schedule of reinforcement (SOR) acquired and maintained cocaine seeking under the control of the conditioned reinforcing properties of the drug-paired CS over a period of three weeks. **(B)** They did not differ in their overall cocaine intake from rats that had been trained to self-administer cocaine under continuous reinforcement (FR1C group). **(C-G)** Analysis of total tissue content of DAT proteins across striatal territories revealed that cocaine exposure, irrespective of schedule of reinforcement, resulted in decreased DAT protein levels across the five striatal territories investigated, i.e., the NacC, the aDLS, the pDLS, the aDMS and the pDMS. some decreases were more specifically pronounced in SOR or FR1C individuals. FR1C rats displayed lower DAT protein levels than controls in the aDMS, in contrast, SOR rats displayed lower DAT protein levels than controls in the aDLS. Inserts show illustrative western blots. (* in **A**: p<0.05 as compared to FI15 block; # in **C-G**: planned comparisons CTL vs cocaine experienced groups, p<0.05; * in **C-G**: pairwise comparisons CTL vs FR1C or CTL vs SOR, p<0.05).

**Figure 3:**
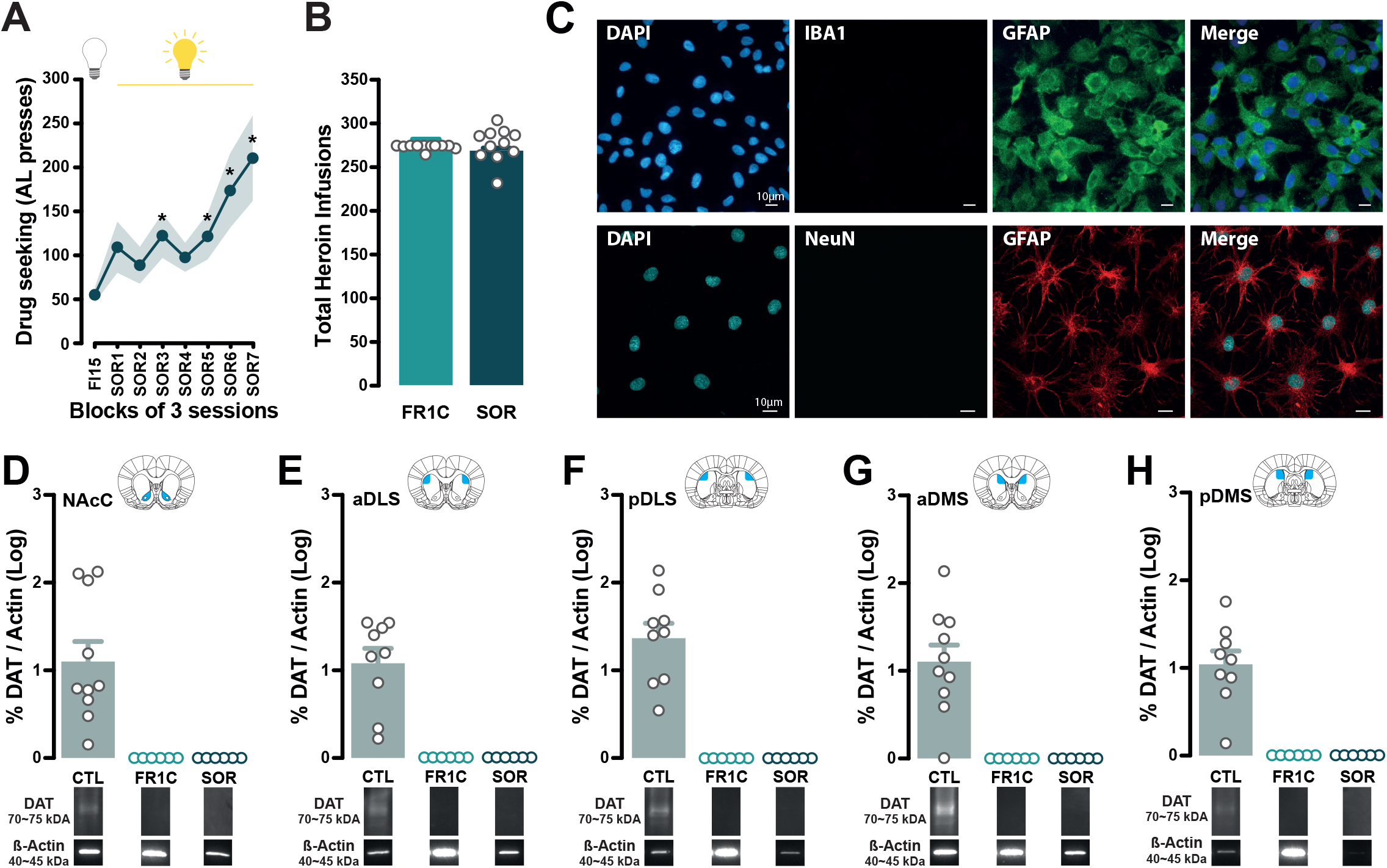
Pan-striatal reductions in astrocytic DAT protein levels are observed following both a history of cocaine taking and cue-controlled cocaine seeking. **(A)** rats trained to seek cocaine under a second-order schedule of reinforcement (SOR) readily acquired and maintained cocaine seeking under the control of the conditioned reinforcing properties of the drug-paired CS over a period of three weeks. **(B)** They did not differ in their overall cocaine intake from rats that had been trained to self-administer cocaine under continuous reinforcement (FR1C group). **(C)** Immunohistochemical analysis of the cultures from astrocytes harvested from 5 striatal territories of each individual brain, revealed pure striatal astrocyte monocultures without microglial or neuronal contamination, as evidenced by the presence of GFAP-positive cells and the absence of Iba-1-positive (top panel) and NeuN-positive cells (bottom panel). **(D-H)** Striatal astrocytes cultured from the striatal territories of control individuals expressed detectable levels of DAT protein, whereas no DAT protein was detected in astrocytes cultured from the FR1C or SOR groups. Inserts show illustrative Western blots. (* in **A**: p<0.05 as compared to FI15 block; * in **D-H**: p<0.05).

### Quantitative polymerase chain reaction (qPCR)

Tissue collection, RNA extraction, reverse transcription, qPCR reaction and analysis were carried out as previously described (Hynes *et al*., 2024). The following primers were used to assess the relative level of DAT mRNA (Quiagen Ref. PPR44664C; Slc6a3; Rn.10093) in comparison to that of Cyclophilin A (Quiagen Ref. PPR06504A; Ppia; Rn.1463) used as housekeeping gene.

### Data and Statistical analyses

Data are presented as means ± SEM and individual data points. Assumptions for normal distribution, homogeneity of variance and sphericity were verified using the Shapiro–Wilk, Levene, and Mauchly sphericity tests, respectively. When assumptions were violated, data were Log transformed.

Differences in Western blot data, qPCR data, and cocaine infusions received were analysed using one-way analysis of variance (ANOVA) with groups (CTL, FR1C, SOR, HC and NC) as between-subject factor. Behavioural data were subjected to repeated measures ANOVAs with sessions as within-subject factor, and compulsivity groups (HC vs NC) as between-subject factor for figure 4. The effect of cocaine exposure on striatal DAT protein levels was assessed by panned comparisons (CTL vs cocaine experienced groups, i.e., FR1C and SOR). Significant main effects were further analysed with Newman-Keuls post-hoc tests. Data from western blot of cultured astrocytes were subjected to a one-sample t-test to test if the control group was significantly greater than zero. Because, as expected, DAT protein levels displayed by naïve rats and those trained to self-administer food did not statistically differ [main effect of group: NacC, aDLS, pDLS, aDMS: all Fs<1; pDMS: F_1,8_ = 1.25, p>0.05], they were pooled to constitute the control (CTL) group.

**Figure 4:**
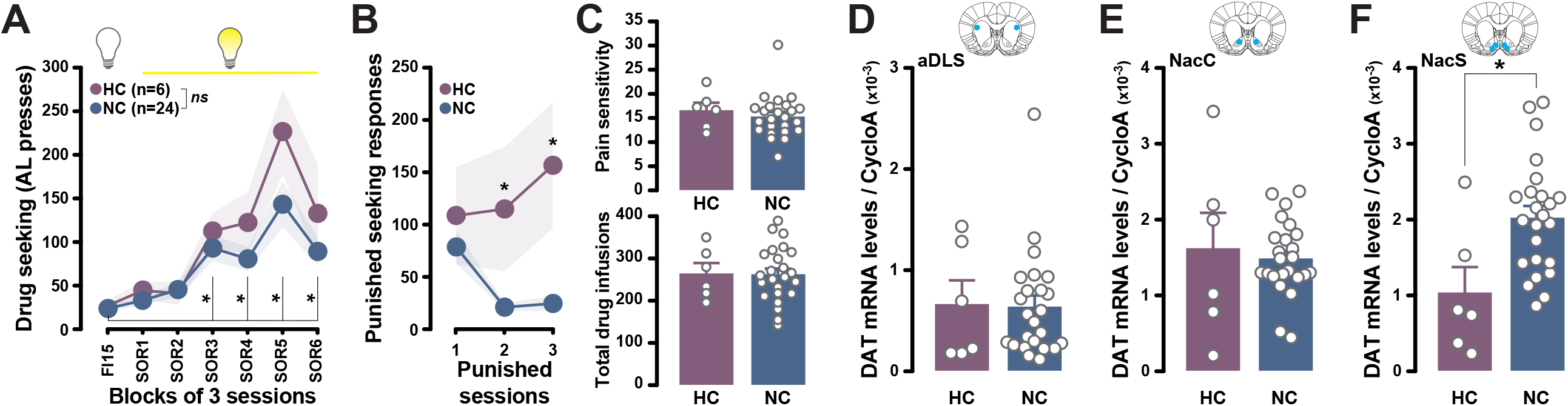
Compulsive incentive cocaine-seeking habits are associated with lower astrocytic DAT mRNA levels in the shell of the nucleus accumbens. **(A)** Rats readily acquired and maintained cocaine seeking under the control of the conditioned reinforcing properties of the drug-paired CS before being challenged over three sessions for their persistence of their incentive seeking habits in the face of punishment. Rats that persisted in their incentive cocaine-seeking habit in the face of punishment (HC rats) did not differ from those that rapidly abstained from seeking over the 3 punished sessions (NC rats). **(B)** HC rats maintained their cocaine seeking across the punished SOR sessions, whereas NC rats dramatically decreased their level of responding in the face of punishment. **(C)** This difference was not attributable to differential pain sensitivity (top panel) or exposure to cocaine throughout self-administration training (bottom panel). HC did not differ from NC rats in the astrocytic DAT mRNA levels measured in their incentive habit striatal network, namely aDLS **(D)** and NacC **(E)**. In marked contrast, HC rats displayed much lower DAT mRNA levels in the NacS than NC rats **(F)**. (* : p<0.05).

For all analyses, significance was set at α = 0.05 and effect sizes reported as partial eta squared (η_p_^2^).

## Results

### Cocaine exposure results in the downregulation of DAT protein levels across the striatum

SOR rats (**Figure 1**) acquired and maintained cocaine seeking under the control of the conditioned reinforcing properties of the drug-paired CS [main effect of session: F_7,63_ = 5.09, p < 0.0001, η_p_^2^ = 0.32] (**Figure 2A**), in line with previous reports (Belin & Everitt, 2008; Puaud *et al*., 2020; Fouyssac *et al*., 2021a). SOR rats did not differ in their overall cocaine intake from FR1C rats [main effect of group: F_1,14_ <1], which had been trained to self-administer cocaine under continuous reinforcement (**Figure 2B**).

Analysis of total tissue content of DAT proteins across striatal territories revealed that cocaine exposure, irrespective of schedule of reinforcement, resulted in decreased DAT protein levels across the five striatal territories investigated, i.e., the NacC, the aDLS, the pDLS, the aDMS and the pDMS [main effect of group: NacC: F_2,23_ = 6.17, p < 0.01, η_p_^2^ = 0.35; aDLS: F_2,23_ = 3.74, p < 0.05, η_p_^2^ = 0.25 ; pDLS: F_2,23_ = 7.83, p < 0.01, η_p_^2^ = 0.40 ; aDMS: F_2,23_ = 4.27, p < 0.05, η_p_^2^ = 0.27; pDMS: F_2,23_ = 5.04, p < 0.05, η_p_^2^ = 0.30] (**Figure 2 C-G**). Planned comparisons confirmed that drug-exposed groups showed lower DAT protein levels than CTL group in each striatal territory even though, pairwise comparisons revealed that some decreases were specifically more pronounced in SOR or FR1C individuals. FR1C rats displayed lower DAT protein levels than controls in the aDMS, in contrast, SOR rats displayed lower DAT protein levels than controls in the aDLS.

The overall decrease in striatal DAT content reflects alterations in both striatal astrocytes and presynaptic dopamine neurons. We further investigated astrocyte-specific alterations in DAT protein levels.

### Striatal astrocytic DAT protein content is profoundly reduced by exposure to cocaine

In a second experiment, SOR rats readily acquired and maintained cocaine seeking under the control of the conditioned reinforcing properties of the drug-paired CS as did those of experiment 1a [main effect of session: F_7,70_ = 5.92, p < 0.0001, η_p_^2^ = 0.37] (**Figure 3A**). SOR rats did not differ in their overall cocaine intake compared to FR1C rats [main effect of group: F_1,14_ =1.48, p >0.05] (**Figure 3B**).

Immunohistochemical analysis of the cultures revealed the presence of pure astrocytic monocultures free from microglial or neuronal contamination, as evidenced by the absence of Iba-1-positive cells (**Figure 3C, top panel**) and NeuN-positive cells (**Figure 3C, bottom panel**).

Cultured astrocytes from CTL rats showed significantly non-zero levels of DAT protein in all striatal territories [NAcC: t_9_ = 4.82, p < 0.001, η_p_^2^ = 0.72; aDLS: t_8_ = 6.39, p < 0.001, η_p_^2^ = 0.83; pDLS: t_8_ = 7.94, p < 0.0001, η_p_^2^ = 0.89; aDMS: t_9_ = 5.84, p < 0.001, η_p_^2^ = 0.79; pDMS: t_8_ = 6.80, p < 0.001, η_p_^2^ = 0.85] (**Figure 3 D-H**). In contrast, no DAT protein could be detected in striatal astrocytes cultured from FR1C or SOR individuals, thereby demonstrating a pan-striatal effect of cocaine exposure on astrocytic DAT protein expression, as we have previously demonstrated to be the case after exposure to heroin (Hynes *et al*., 2024).

Having previously shown that astrocytic DAT mRNA can be detected using qPCR even under conditions, such as after heroin exposure, resulting in astrocytic protein levels low enough not to be detectable using western blot (Hynes *et al*., 2024), we used qPCR to assess whether compulsive incentive cocaine-seeking habits were associated with further alterations in post-synaptic striatal, astrocytic, DAT mRNA levels in SOR-trained individuals (Hynes *et al*., 2024). Considering the recent body of evidence implicating monoaminergic innervation of the Nac Shell (NAcS) in compulsive behaviours (Chernoff *et al*., 2024), we measured DAT mRNA levels from this striatal region as well as the striatal incentive habit circuitry, namely NacC and aDLS, in rats with a compulsive incentive cocaine-seeking habit (High Compulsive, HC) or not (Non-Compulsive, NC).

### Compulsive incentive cocaine-seeking habits are associated with decreased DAT mRNA in the NacS

As in the previous two experiments, rats readily acquired and maintained cocaine seeking under the control of the conditioned reinforcing properties of the drug-paired CS [main effect of session: F_6,168_ = 23.62, p < 0.0001, η_p_^2^ = 0.45] (**Figure 4A**) before being challenged over three sessions for their persistence of their incentive seeking habits in the face of punishment. Unbiased k-means clustering on the number of shocks rats received during drug-free seeking periods revealed two very different populations. HC rats maintained cocaine seeking across the punished SOR sessions while NC rats dramatically decreased their level of responding in the face of punishment [main effect of group: F_1,28_ = 10.78, p < 0.01, η_p_^2^ = 0.28; group x session interaction: F_2,56_ = 3.93, p < 0.05, η_p_^2^ = 0.12] (**Figure 4B**). This difference was neither attributable to differential pain sensitivity nor exposure to cocaine throughout self-administration training [all Fs < 1] (**Figure 4C**) or acquisition and maintenance of cue-controlled cocaine seeking prior to be exposed to contingent punishments [main effect of group: F_1,28_ = 1.22, p > 0.1; group x session interaction: F_6,168_ = 1.74, p > 0.1] (**Figure 4A**).

DAT mRNA levels did not differ between HC and NC rats in the striatal incentive habits network, namely aDLS and NacC [all Fs < 1] (**Figure 4D and 4E**). However, HC rats displayed lower DAT mRNA levels than NC rats in the NacS [F_1,28_ = 8.22, p < 0.01, η_p_^2^ = 0.23] (**Figure 4F**).

## Discussion

Exposure to cocaine, either under continuous reinforcement or SOR (promoting aDLS dopamine-dependent incentive habits), resulted in a profound decrease in DAT protein and mRNA levels in the striatum that was mostly attributable to alterations within astrocytes across the ventral and dorsal territories of the striatum.

These results parallel previous reports of a reduction in DAT binding sites in the caudate, putamen and Nac of humans with cocaine addiction (Hurd & Herkenham, 1993). Similarly, studies in non-human primates have shown a decrease in DAT binding sites, initially confined to the ventral striatum, that progressively spreads to more lateral and dorsal areas of the striatum over the time course of continued cocaine use (Letchworth *et al*., 2001).

Drug-induced pan-striatal reduction in astrocytic DAT protein levels to levels undetectable by western blot assays observed here following cocaine exposure is not unprecedented. A similar drug-induced decrease in astrocytic DAT protein expression was also observed after heroin exposure (Hynes *et al*., 2024), suggesting that such alterations in astrocytes across the striatum are triggered by drug-induced increase in dopamine release across drug classes [i.e., DAT blockade by cocaine (Ritz *et al*., 1987) and disinhibition of midbrain dopaminergic neurons by opioids (Johnson & North, 1992)]. In astrocyte monocultures from drug naive rats, we have previously shown that chronic exposure to dopamine, at levels equivalent to those induced by intravenous cocaine infusions, results in decreased DAT protein levels in some striatal regions (Hynes *et al*., 2024). Together, these data suggest that prolonged exposure to high dopamine levels, as experienced following cocaine, or heroin self-administration, may result in a downregulation of astrocytic DAT protein levels.

The present cocaine-induced alterations in astrocytic DAT were observed in the incentive habit striatal circuit, namely NacC and aDLS, even after a short period of training under continuous reinforcement for cocaine but not for food. Together with our recent demonstration of heroin-induced decrease in striatal astrocyte DAT protein and mRNA levels, this observation supports our hypothesis that exposure to drugs of abuse triggers downregulation of dorsal striatal astrocytic DAT protein levels before the development of incentive drug-seeking habits.

While a history of cocaine self-administration, irrespective of the schedule of reinforcement triggers a decrease in total tissue DAT protein levels (both neuronal and astrocytic), we previously showed that exposure to heroin downregulated total tissue DAT exclusively in the aDLS of rats trained under SOR and even upregulated DAT in the aDLS and pDLS of rats trained to self-administer heroin under continuous reinforcement. This suggests that while a common mechanism between addictive drugs likely underlies astrocytic adaptations in DAT expression, drug-specific processes may elicit distinct, potentially compensatory, neuronal adaptations in DAT expression. Although further research is required, it may be speculated that such differences in neuronal alterations following cocaine or heroin exposure could stem from differences in the characteristics of the hyperdopaminergic states induced by these two drugs, direct drug interactions with astrocytic targets or even distinct indirect mechanisms at the system-level. Indeed, chronic *in vitro* opiate application downregulates DAT protein in cultured striatal astrocytes (Hynes *et al*., 2024), indicating a role for astrocytic opioid receptors in heroin-induced downregulation of DAT protein. Interestingly, cocaine itself directly interacts with DAT, causing internalisation of the transporter in cultured neurons (Saenz *et al*., 2023). Whether this also occurs in astrocytes remains to be established.

While drug self-administration and incentive drug-seeking habits are associated with a decrease in DAT protein and mRNA levels across the striatum, compulsive incentive cocaine-seeking habits are characterised by a decrease in astrocytic DAT mRNA levels in the NacS, but not in the incentive habit striatal system. Rats that persisted in their incentive cocaine seeking habit (HC rats) did not differ from those whose responding was profoundly reduced by the same electric foot shocks in their levels of DAT mRNA in the incentive habits-associated striatal system (NacC and aDLS). This suggests that the inability to disengage aDLS dopamine-dependent incentive habits in the face of punishment, as we have previously shown to be the case for compulsive alcohol seeking (Giuliano *et al*., 2019), is not directly related to astrocytic DAT levels in this system.

HC rats displayed lower DAT mRNA levels in the NacS than NC rats. This observation is in agreement with the role of NacS monoaminergic mechanisms have been shown to play in several compulsive behaviours (Sturm *et al*., 2003; Chernoff *et al*., 2024). The NacS is the only striatal territory innervated by noradrenergic neurons from the Locus Coeruleus (Berridge *et al*., 1997; Delfs *et al*., 1998). DAT can clear extracellular noradrenaline, albeit with less efficacy than it does dopamine (Ranjbar-Slamloo & Fazlali, 2019), and cocaine has a relatively high affinity for the norepinephrine transporter (Koe, 1976), which also reuptakes dopamine (Moron *et al*., 2002). The present findings therefore suggest that compulsive incentive cocaine-seeking habits, the vulnerability to which is predicted by impulsivity- and stickiness-related structural and functional alterations in several parallel loops of the corticostriatal circuitry, including the ventral striatum (Jones *et al*., 2024), may also result from drug-induced hyper dopaminergic and noradrenergic states in the NacS.

In summary, the results of the present study demonstrate that exposure to cocaine self-administration triggers similar pan-striatal adaptations in astrocytic DAT to those produced by exposure to heroin while the compulsive expression of incentive cocaine seeking habits is associated with a selective decrease in DAT mRNA in the NacS astrocytes. These findings suggest that alterations in astrocytic DAT may represent a common mechanism underlying the striatal dopaminergic adaptations that lead to the development of compulsive incentive drug-seeking habits across addictive drugs.

## Acknowledgements

This work was supported by a UKRI MRC programme grant to Barry J Everitt, DB, Amy Milton, Trevor W Robbins and Jeffrey W Dalley (MR/N02530X/1) and a UKRI MRC research grant to DB (MR/W019647/1). MF was supported by a Leverhulme Trust Early Career Fellowship (ECF-2021-451) and the Isaac Newton Trust (1.08(n)). TH was supported by a Leverhulme Trust Early Career Fellowship (ECF-2024-242) and the Isaac Newton Trust (24.08).

## Author contributions

MF and DB conceptualised and designed the experiments. MF carried out the behavioural procedures as well as western blot assays. ABR performed the qPCR assays. MF, TH and DJ performed the primary astrocyte cultures and immunohistochemistry assays. MF and DB wrote the manuscript.

## Data availability

Data will be made available on the Cambridge repository after acceptance of this manuscript for publication.

