## Supplemental Figure S1 for "Compulsion derived from incentive cocaine-seeking habits is associated with a downregulation of the dopamine transporter in striatal astrocytes"

**This PDF file includes:**

Figures S1

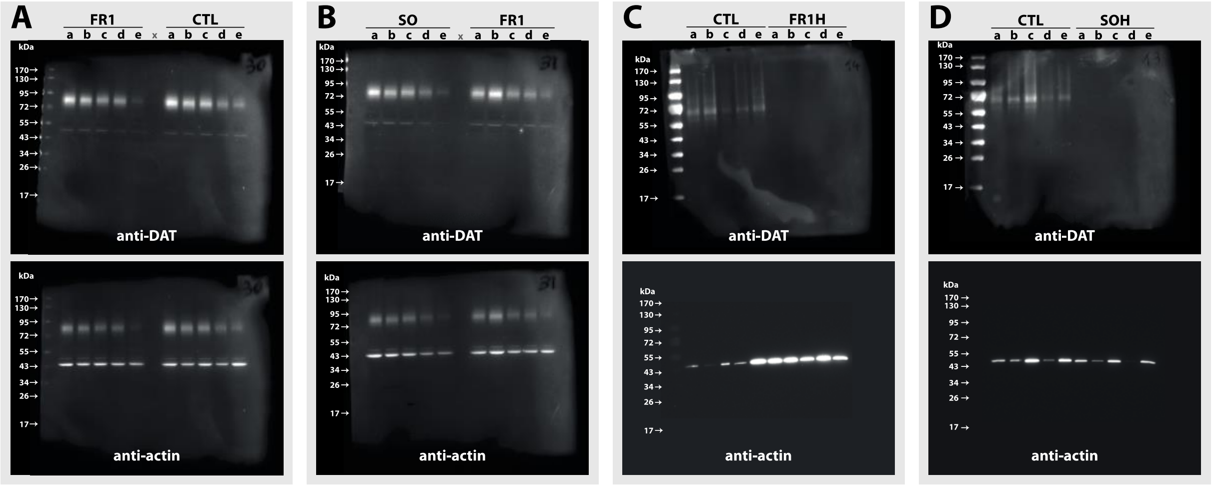


a: aDLS (anterior dorsolateral striatum); b: pDLS (posterior dorsolateral striatum); c: aDMS (anterior dorsomedial striatum); d: pDMS (posterior dorsomedial striatum); e: NacC (nucleus accumbens core)

**Figure S1. Raw uncropped western blot images.**
**A - B)** complete unedited western blot images related to data presented in **Figure 2** from micro-punched samples pertaining to the three experimental conditions CTL, FR1C and SOR. **C - D)** complete unedited western blot images related to data presented in **Figure 3** from cultured astrocytes samples pertaining to the three experimental conditions CTL, FR1C and SOR. DAT proteins are revealed at ~70/75 kDa (top panels) and actin at ~40/45 kDa (bottoms panels).
